# Dishevelled controls bulk cadherin dynamics and the stability of individual cadherin clusters during convergent extension

**DOI:** 10.1101/2022.06.06.495004

**Authors:** Robert J. Huebner, John B. Wallingford

## Abstract

Animals are shaped through the movement of large cellular collectives. Such morphogenetic processes require cadherin-based cell adhesion to maintain tissue cohesion and planar cell polarity to coordinate movement. Despite a vast literature surrounding cadherin-based adhesion and planar cell polarity it is unclear how these molecular networks interface. Here we investigate the relationship between cadherins and planar cell polarity during gastrulation cell movements in *Xenopus laevis*. We first assessed the bulk cadherin localization and found that cadherins were enriched at a specific subset of morphogenetically active cell-cell junctions. We then found that disruption of planar cell polarity desynchronized temporal cadherin and actin dynamics. Next, using super-resolution time-lapse miscopy and quantitative image analysis we were able to measure the lifespan and size of individual cadherin clusters. Finally, we show that planar cell polarity not only controls the size of cadherin clusters but, more interestingly, regulates cluster stability. These results reveal an intriguing link between two essential cellular properties, adhesion and planar polarity, and provide insight into the molecular control of morphogenetic cell movements.

**Highlights:** Planar cell polarity is required for cadherin spatiotemporal dynamics.

Super-resolution imaging reveals the spatiotemporal patterns of cadherin clustering.

Cadherin cluster lifespan and size are highly heterogenous.

Planar cell polarity controls cadherin cluster stability and size.

## Introduction

Embryonic development requires the stereotyped movement of cellular collectives. An animal’s anterior-posterior (head-to-tail) axis is established in part through a specific form of collective cell movements termed convergent extension (CE) in which cells converge along one axis resulting in extension along the perpendicular axis (Keller and Sutherland, 2020). CE is a deeply conserved and essential cellular process and has been described in organisms ranging from nematodes to mammals (Huebner and Wallingford, 2018), and failure of CE is linked to human birth defects (Wallingford et al., 2013). Thus, cultivating a deeper knowledge of CE will not only inform our understanding of this essential developmental process but will also provide insight into the etiology of human disease.

Vertebrate CE is patterned through the asymmetric localization of planar cell polarity (PCP) proteins, which allows cells to sense directionality along the anterior-posterior axis (Butler and Wallingford, 2017). CE also requires cell-cell adhesion to maintain tissue integrity and to transduce forces between neighboring cells (Lecuit and Yap, 2015). Such adhesion is mediated by cadherins, homophilic cell adhesion molecules that are required for a multitude of collective cell behaviors (Arslan et al., 2021) including vertebrate CE (Lee and Gumbiner, 1995). While PCP and cadherin-based cell adhesions have been studied to great depth, relatively little is known about how these cellular activities interface during CE.

PCP signaling has been shown to function upstream of cadherin mediated cell adhesion in *Xenopus, Drosophila*, zebrafish, mice, and cell culture (Dohn et al., 2013; Kraft et al., 2012; Mirkovic et al., 2011; Nagaoka et al., 2014; Tatin et al., 2013) and multiple mechanisms have been proposed for the underlying interaction. One such mechanism is that PCP controls the ability of cadherins to form higher ordered clusters that modulate adhesion (Kraft et al., 2012). Recently we found that perturbation of PCP disrupted cis-clustering of *Xenopus* Cdh3 (aka C-cadherin, aka mammalian P-cadherin) (Huebner et al., 2021). This result led us to perform a deeper investigation into the relationship between PCP and cadherin during axis elongation in *Xenopus*.

Here, we show that Cdh3 is enriched at specific cell-cell junctions during *Xenopus* CE. Next, we determine that PCP was required for the temporal coordination of cadherin and actin dynamics. We then developed a new quantitative imaging approach to measure the size and, importantly, lifespan of individual cadherin clusters, revealing that clusters are highly dynamic and heterogeneous in both size and lifespan. Finally, we demonstrate that perturbation of PCP reduced cluster size and more interestingly reduced cluster lifespan. These results confirm a relationship between PCP and cadherin clustering and specifically show that PCP is required for stabilizing clusters. These data not only improve our understanding of two essential molecular networks, PCP and cadherin-based adhesions, but also deepens our understanding of the molecular control of a fundamental developmental process, convergent extension.

## Results

### Cadherins are enriched at shortening cell-cell junctions during CE

*Xenopus* CE results from cell intercalation, during which cells come together along the mediolateral axis and are pushed apart along the anterior-posterior direction (Shih and Keller, 1992) (Fig.1A). During this process, mediolaterally-oriented junctions (termed “v-junctions” by convention) shorten and are replaced by anterior-posterior oriented junctions (termed t-junctions) (Bertet et al., 2004; Blankenship et al., 2006) (Fig.1A). Cadherins are planar polarized to t-junctions during CE in the *Drosophila* germ band (Blankenship et al., 2006), so we began our study by asking if cadherins are similarly polarized during CE in *Xenopus*.

**Figure 1:**
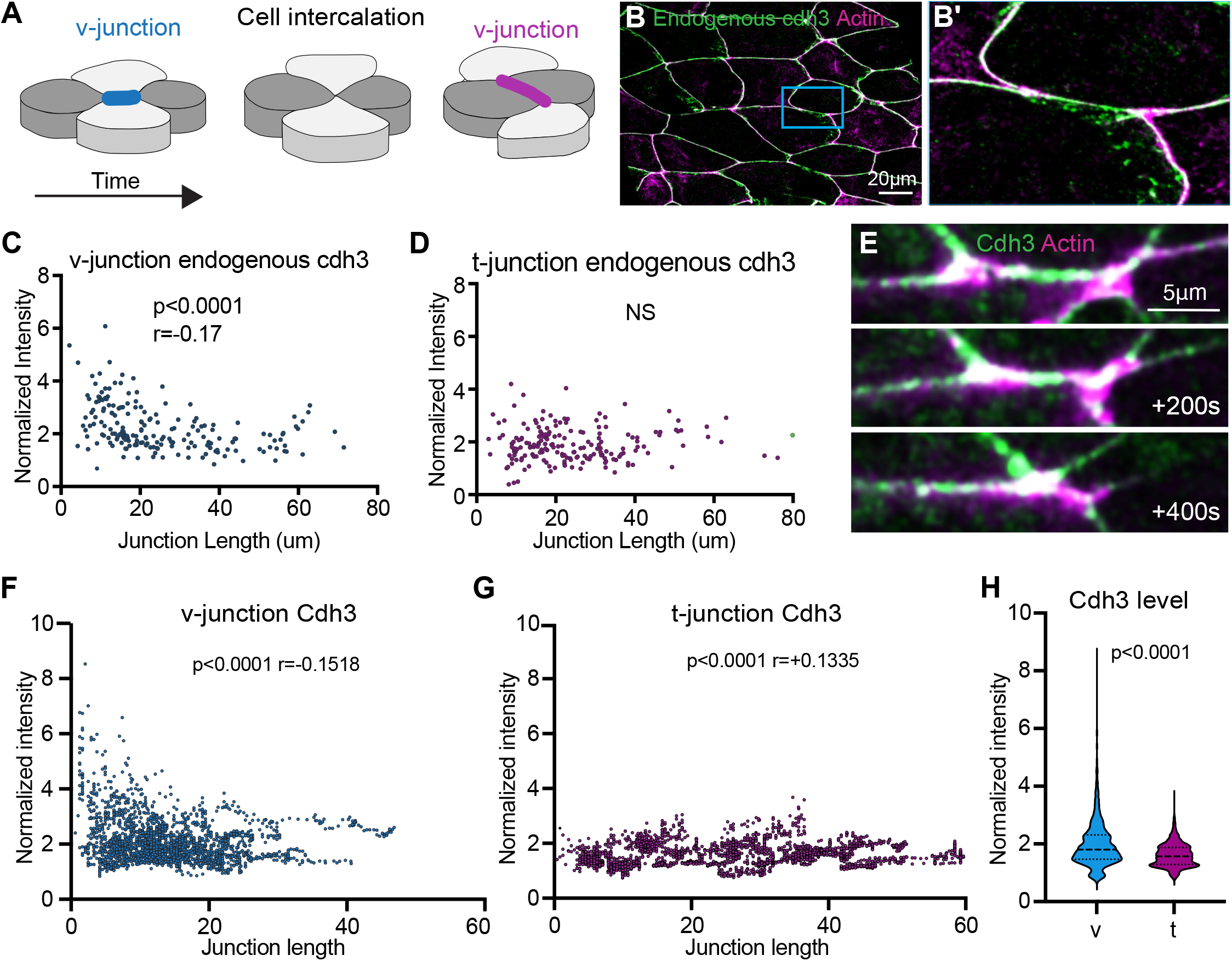
Cdh3 is enriched at shortening v-junctions. **A**. Schematic depicting the cell intercalation movements that drive convergent extension. **B**. Single image from a time lapse movie of *Xenopus* mesodermal cells undergoing convergent extension. Cells are labeled for Cdh3 (Cdh3-GFP) in green and actin (lifeact-RFP) in magenta. **B’**. Time series of images from the inset in Fig.1B. Here we are zooming in to highlight a single shrinking v-junction. **C**. Graph of the mean intensity at shortening v-junctions or at t-junctions. Intensities were collected from time lapse movies and were normalized to account for bleaching. Conditions were statistically compared using a Mann Whitney test. **D**. Graph comparing mean Cdh3 intensity to v-junction length. The relationship between Cdh3 intensity and v-junction length was assessed by Spearman correlation. **E**. Comparison of Cdh3 intensity to t-junction length. Spearman correlation was used to assess the relationship between Cdh3 intensity and t-junction length. **F**. Still image from a time lapse movie of *Xenopus* mesodermal cells expressing dominant negative dishevelled-2 (Xdd1). Xdd1 specifically disrupts PCP signaling and inhibits convergent extension cell behaviors. Here cells are labeled for Cdh3 in green and actin in magenta. **G**. Graph comparing Cdh3 intensity at shortening v-junctions or cell junctions where Xdd1 is present in the background. Conditions were statistically compared using a Mann Whitney test. **H**. Comparison of Cdh3 intensity to junction length from cells expressing Xdd1. The relationship between intensity and junction length was assessed by Spearman correlation.

To assess Cdh3 polarity we used immuno-staining to observe the endogenous Cdh3 localization at v-and t-junctions. Visual inspection showed that Cdh3 was present at v-junctions (Fig.1B-B’). We next measured endogenous Cdh3 fluorescent intensities at v-and t-junctions. Interestingly we observed a slight but significant negative correlation between Cdh3 intensity and v-junction length, such that Cdh3 was enriched at the shortest v-junctions (Fig.1C). No such correlation was observed at t-junctions (Fig.1D). This result led us to ask how Cdh3 levels change specifically at shortening v-junctions.

To address this, we used a well-characterized, functional Cdh3-GFP fusion protein (Huebner et al., 2021; Pfister et al., 2016) to visualize and quantify Cdh3 protein localization in real-time at shortening v-junctions (Fig.1E). Quantification of mean Cdh3-GFP intensity at shortening v-junctions showed a clear enrichment of Cdh3 during junction shortening (Fig.1F). No such correlation was observed between junction length and Cdh3 intensity at t-junctions (Fig.1G). Further, when comparing Cdh3 intensity at shortening v-junctions to t-junctions there was an enrichment for Cdh3 at the shortening v-junctions (Fig.1H). Our interpretation of this data is that Cdh3 is not necessarily polarized during Xenopus CE but instead becomes increasingly enriched at shortening v-junctions.

These results are intriguing as cadherins in *Xenopus* CE are not planar polarized as observed in *Drosophila*, which has been the paradigm for studying cadherin function during CE (Blankenship et al., 2006). These results are further surprising because the patterns of actomyosin localization are largely the same in these two systems (Bertet et al., 2004; Blankenship et al., 2006; Butler and Wallingford, 2018; Shindo and Wallingford, 2014). Thus, it seems there is a fundamental difference in Cdh3 function during *Xenopus* mesenchymal CE compared to cadherins during Drosophila germ band extension CE.

### Planar cell polarity is required for synchronized cadherin and actin oscillations

The actomyosin cytoskeleton provides the force to move cellular collectives and it has become increasingly clear that actin and myosin function through oscillatory periods of activity and inactivity (Coravos et al., 2017). In *Xenopus*, actomyosin oscillations are observed at v-junctions and proper tuning of the frequency and amplitude of oscillations is required for efficient cell intercalation (Huebner et al., 2022; Shindo et al., 2019). Further, Cdh3 has also been observed to oscillate at v-junctions in *Xenopus* and oscillatory changes in cadherin cluster size correlate with v-junction shortening events (Huebner et al., 2021). These results led us to ask if there is a temporal relationship between Cdh3 and actin oscillations.

To this end, we collected time lapse movies of Cdh3 and actin at shortening v-junctions. We observed oscillatory pulses of Cdh3 and actin and these were clearly correlated in time (Fig.2A-A’’, B). Moreover, cross-correlation analysis confirmed a strong positive correlation between actin and Cdh3 oscillations (Fig.2C). Interestingly, such coordinated behavior was specific to shortening v-junctions, as Cdh3 and actin intensity were poorly coordinated at t-junctions when compared to v-junctions (Fig.2D-E). These data show a clear coordination of Cdh3 and actin oscillations specifically at shortening v-junctions which led us to ask if PCP was required for this asymmetric oscillatory behavior.

**Figure 2:**
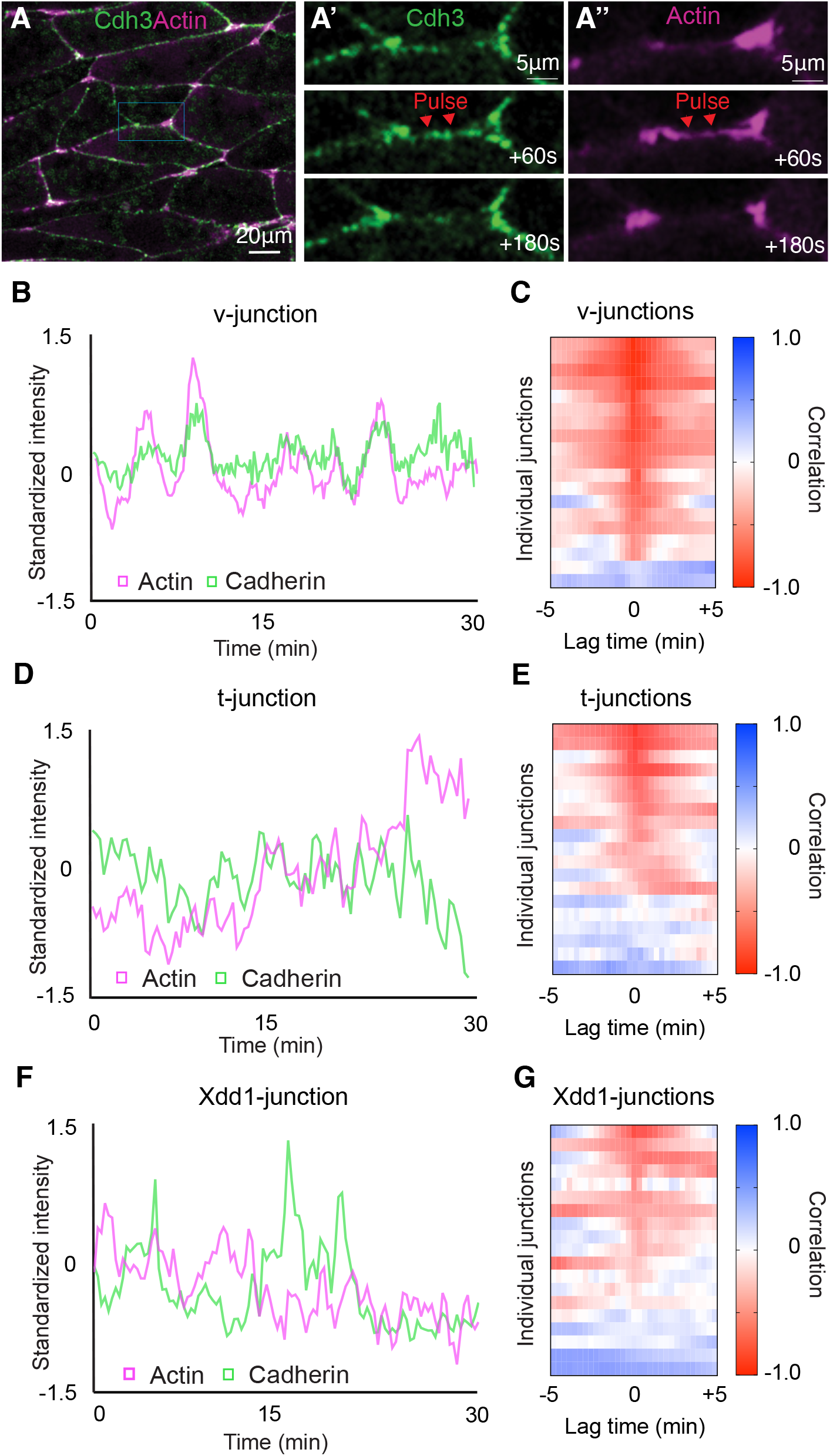
PCP is required for synchronized cadherin and actin dynamics. Frame from a time lapse movie of intercalating cells labeled for Cdh3 in green and actin in magenta. **A’**. Time series of Cdh3 from the inset in Fig.2A. Here we are highlighting a shortening v-junction and showing that Cdh3 undergoes oscillatory pulses at said junctions. **A’’**. Time series of actin from the inset in Fig.2A. In this time series there is an actin pulse that occurs concurrently with the Cdh3 pulse shown in Fig.2A’. **B**. Graph plotting Cdh3 and actin intensity over time at a shortening v-junction. Here we observe concurrent peaks of Cdh3 and actin. Intensities were normalized to account for bleaching and then standardized to emphasize the overlap in the two signals over time. **C**. Heatmap of the cross-correlation analysis for 19 shortening v-junctions. Each line on the y-axis of this heatmap represents the cross-correlation for a single shortening v-junction. The value of the correlation is represented in color code with red being positive correlation and blue being negative correlation. The x-axis shows the lag time and the fact we observe a high positive correlation at lag time 0 indicates that Cdh3 and actin pulses occur concurrently. **D**. Graph plotting Cdh3 and actin intensities at t-junctions. **E**. Heatmap of the cross-correlation analysis for 19 t-junctions. **F**. Graph of Cdh3 and actin over time measured at junctions where Xdd1 was present in the background. **G**. Heatmap of the cross-correlation analysis for 19 junctions with Xdd1 in the background.

Next, we used a dominant negative dishevelled-2 (Xdd1), that specifically disrupts PCP, to test if PCP was required for the coordinated Cdh3 and actin oscillations (Sokol, 1996; Wallingford et al., 2000). We collected time lapse movies of Cdh3 and actin with Xdd1 in the background and here we specifically chose junctions oriented as v-junctions. We found that Cdh3 and actin were poorly cross correlated after PCP perturbation (Fig.2F-G). These data suggest that PCP has a role in coupling Cdh3 and actin dynamics and overall, these results show that Cdh3 and actin undergo PCP dependent coordinated oscillations specifically at v-junctions.

### Planar cell parity is not required for cadherin turnover

Having determined that PCP was required for the bulk localization and temporal dynamics of Cdh3, we next asked how PCP controls these features. Cadherin protein levels and dynamics can be modulated by altering the protein turnover at cell junctions (Cavey et al., 2008). One method to assess bulk protein turnover is fluorescence recovery after photobleaching (FRAP) (Reits and Neefjes, 2001). We therefore performed FRAP on Cdh3 (Supp Fig.1A) in the presence or absence of Xdd1. Surprisingly, we observed no difference in the FRAP curves, the recovery half-time, or the recovery mobile fraction (Supp Fig.1B-D). These results show that PCP does not govern bulk Cdh3 turnover at cell-cell junctions and instead must regulate some other aspect of Cdh3 behavior.

### Cadherins clusters are dynamic and heterogenous during *Xenopus* CE

Cadherins form intercellular and intracellular interactions with fellow cadherins to generate adhesion complexes, termed clusters, that are observable from the nano to micron scale (Yap et al., 2015). We have recently shown that cadherin clustering is required for CE in *Xenopus* and further that cadherin cluster size and temporal dynamics are important for tuning the local mechanics of cell-cell junctions during CE (Huebner et al., 2021). We also observed that perturbation of PCP altered cadherin cluster sizes (Huebner et al., 2021). Thus, it seems possible that PCP controls bulk Cdh3 dynamics by modulating cadherin clusters. However, our previous work was averaged over large datasets and did not allow us to study the behavior of individual clusters. We therefore sought to develop a method to measure the size and importantly the lifespan of individual Cdh3. It is of note that we are particularly interested in cluster lifespan as temporal clustering dynamics modulates v-junction shortening (Huebner et al., 2021) and because, to our knowledge, there are few or no current methods to measure cluster lifespan.

Here we collected super-resolution time lapse movies of the classic cadherin Cdh3 during CE. We were able to clearly observe the formation and dissipation of submicron scale cadherin clusters (Fig.3A-A’’’). From these movies it was apparent that cluster size and lifespan were heterogeneous (Fig.3A-A’’’). To measure cluster size and lifespan, we first made kymographs across the length of individual cell-cell junctions (Fig.3B-B’). Kymographs were then thresholded to remove background and smoothed along the time axis to make clusters a continuous line in the image (Fig.3B’’). A binary mask was then applied to the kymographs which allowed us to identify each cluster as an individual object and objects touching the edges of the kymograph were discarded (Fig.3B’’’). Using this method, we were able to measure the size, lifespan, and number of clusters observed at a cell junction (Fig.3B’’’-E). Importantly, the mean cluster size observed with this new method matched well with the previously published mean cluster size observed in our large averaged datasets, providing evidence that our new approach is effective (Huebner et al., 2021).

**Figure 3:**
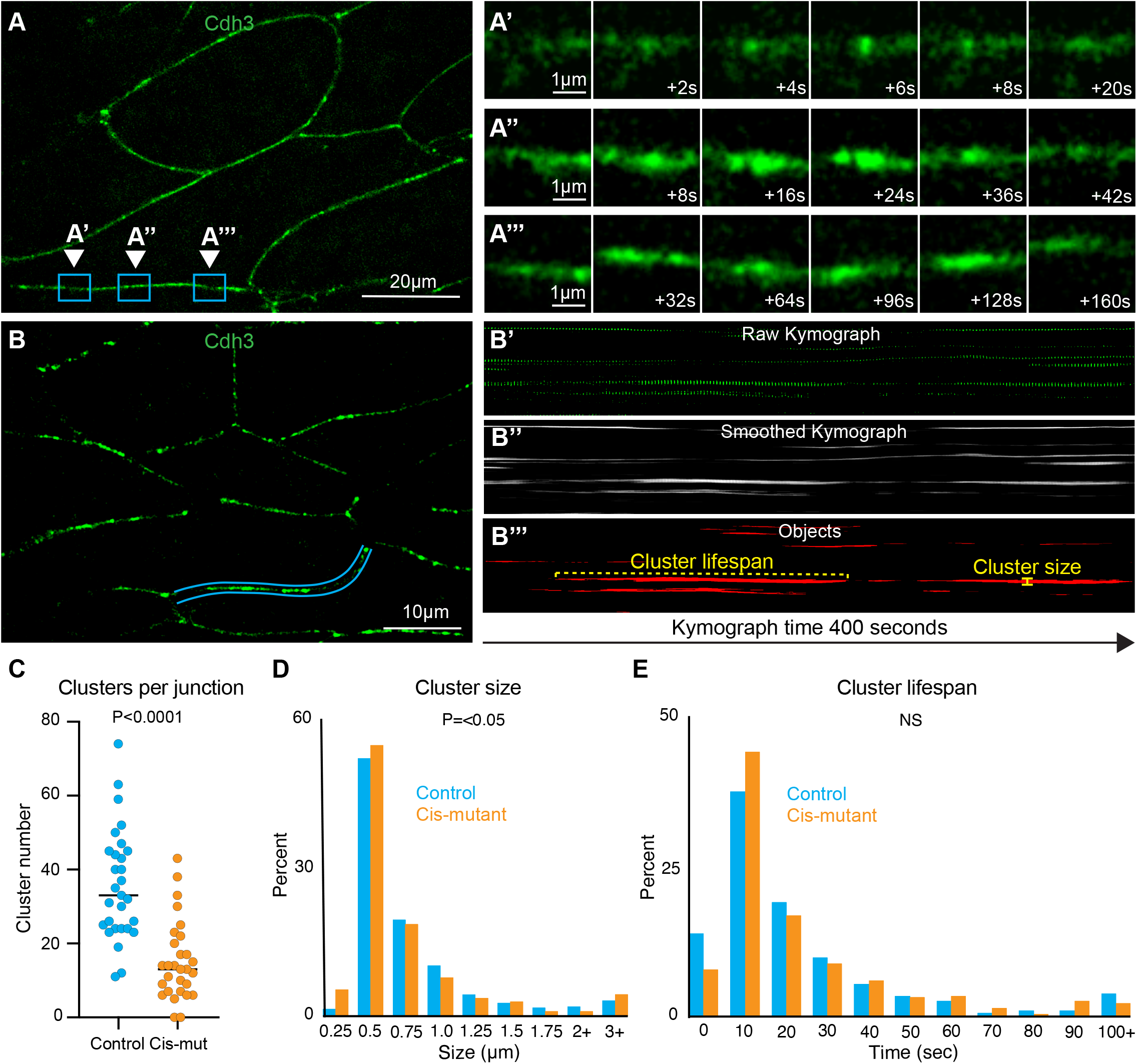
Cadherin clusters display heterogenous spatiotemporal dynamics during CE. **A**. Still frame from a super-resolution time lapse movie of Cdh3. Three different regions of a single junction are highlighted, each region is the position where a cluster formed and dissipated. These regions were chosen to highlife the heterogeneity in cluster size and lifespan. **A’**. Frames from the time lapse movie shown in A with a zoom on the inset labeled A’. These images show a small, short lifespan Cdh3 cluster. **A’’**. Frames from the time lapse movie shown in A with a zoom on the inset labeled A’’. Images show a large cluster with an intermediate lifespan. **A’’’**. Frames from the time lapse movie shown in A with a zoom on the inset labeled A’’’. Images show a particularly long lived Cdh3 cluster. **B**. Still frame from a time lapse super-resolution movie of Cdh3. Here the blue lines highlight the cell junction that was used to make the kymographs in Fig.4B’-B’’’. **B’**. The raw kymograph of the cell junction highlighted in Fig.4B. This kymograph shows Cdh3 over a 400 second time period beginning at the left side of the image. **B’’**. Smoothed image of the kymograph shown in Fig.4B’. This kymograph was smoothed along the time axis to connect the Cdh3 spots into uninterrupted clusters. **B’’’**. This image shows a thresholded version of the kymograph in Fig.4B’’. Here the kymograph was thresholded and converted to a binary image which allowed us to identify clusters as individual objects. Objects that touched the edges of the kymograph were removed so that we only evaluated complete clusters. The cluster lifespan was then determined as the cluster length and the cluster size was determined by the width of each cluster. **C**. Graph showing the number of clusters per junction for control and Cdh3-cis-mutant junctions. Each spot represents a single junction, and all junctions were measured over a 400 second timeframe. Conditions were statistically compared using a Mann Whitney test. **D**. Histogram displaying the size distribution of control and Cdh3-cis-mutant clusters. Conditions were statistically compared using a Kolmogorov-Smirnov test. **E**. Histogram displaying the Cdh3 cluster lifespan distribution for control and Cdh3-cis-mutant clusters. Conditions were statistically compared using a Kolmogorov-Smirnov test.

We next tested the validity of this new method by applying it to two negative controls, a membrane marker that does not form clusters and a well characterized cadherin mutant with disrupted clustering (Cdh3-cis-mutant)(Harrison et al., 2011; Huebner et al., 2021). As expected, we observed almost no clustering with the membrane marker (Supp. Fig.2A-A’’’). To assess clustering in the Cdh3-cis-mutant, we used a previously developed method for knockdown of the wildtype Cdh3 and replacement with the Cdh3-cis-mutant (Huebner et al., 2021). As expected, expression of the Cdh3-cis-mutant resulted in a significant reduction in the number of clusters observed at each cell junction using the new method (Fig.3C) (Supp. Fig.3A-A’’’). Interestingly, however, when we compared the distributions of Cdh3 cluster size and lifespan with the mutant we found only a modest difference in size and no difference in lifespan (Fig.3D, E). These data suggest that the mutant inhibits the ability of clusters to form but has little effect on the dynamics of clusters that do form. Importantly, these results show that we have developed an effective method to measure individual Cdh3 cluster dynamics.

### Planar cell polarity controls individual cadherin cluster size and stability

With this new cluster analysis method, we sought to develop a more refined understanding of how PCP regulates Cdh3 clustering. Here we overexpressed Xdd1 and collected super-resolution time lapse movies of Cdh3 (Fig.4A). When visualizing cell junction kymographs of Xdd1 expressing cells we immediately noticed that the clusters appeared to have much shorter lifespans when compared to control kymographs (Fig.4A’, B). Measurement of the number of clusters per junction showed a significant reduction in clusters in the presence of Xdd1, consistent with our observations of the Cdh3-cis-mutant (Fig.3C). We also observed a reduction in cluster size when comparing with Xdd1 (Fig.4D). However, the most striking result was a reduction in cluster lifespan in the presence of Xdd1 (Fig.4E). This change in cluster lifespan was specific to Xdd1 as it was not observed in the Cdh3-cis-mutant. These results indicate that PCP has a role in stabilizing Cdh3 clusters and provide an exciting new link between PCP and cadherin-based cell adhesions.

**Figure 4:**
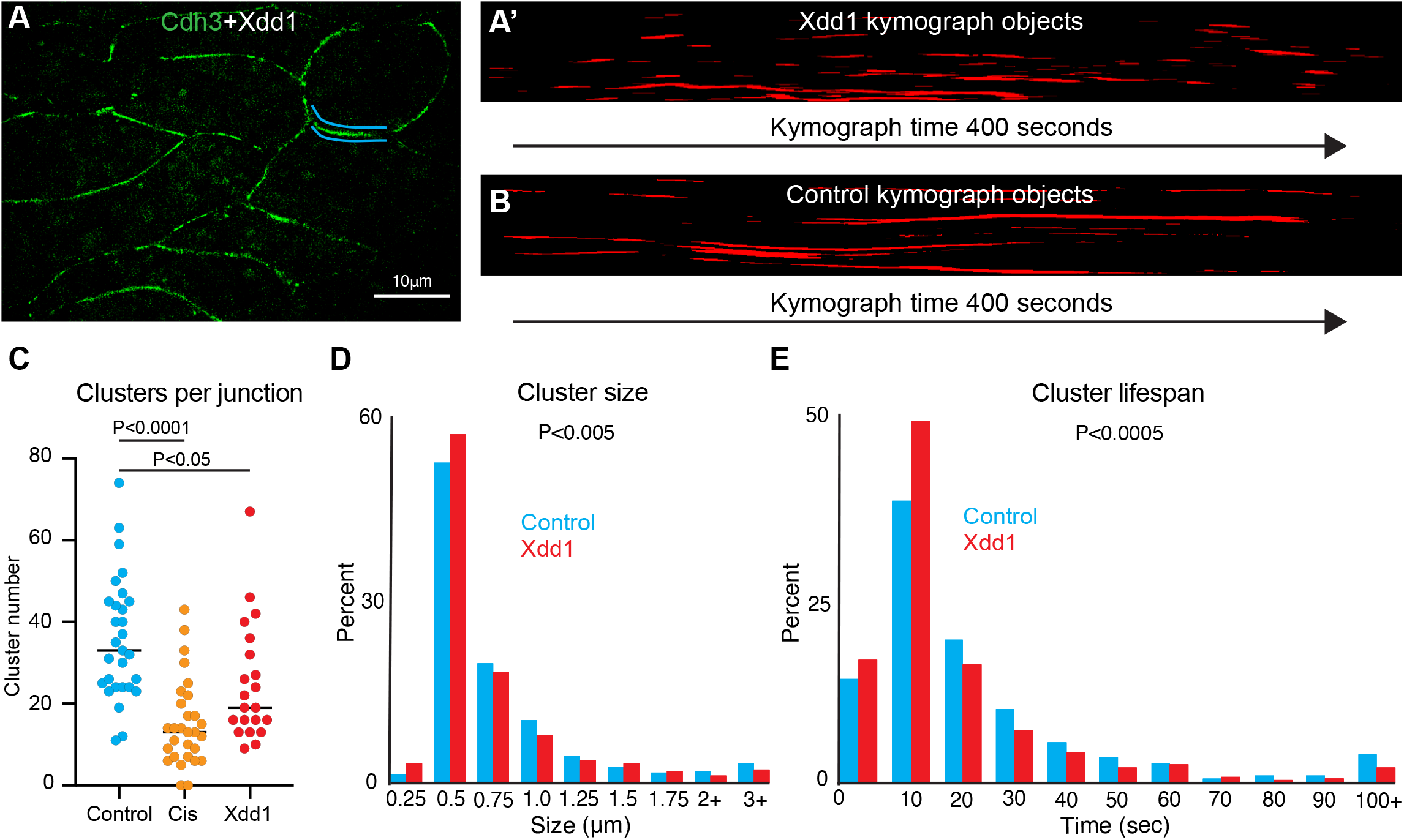
PCP regulates cadherin cluster stability. **A**. Still frame of cells labeled for Cdh3 in green with Xdd1 in the background. Blue lines highlight the junction shown in the kymograph in Fig.5A’. **A’**. Kymograph of the cell junction highlighted in Fig.5A. This kymograph has been smoothed, thresholded, and converted to objects. **B**. Kymograph of a control cell junction for comparison to the Xdd1 junction. This kymograph has also been smoothed, thresholded, and converted to object. **C**. Graph of the number of Cdh3 clusters per junction for control, Cdh3-cis-mutant, and Xdd1. Each dot represents one junction over a 400 second measurement interval. Conditions were compared using an ANOVA. **D**. Histogram displaying the size distribution of control clusters and clusters with Xdd1 in the background. Conditions were statistically compared using a Kolmogorov-Smirnov test. **E**. Histogram displaying the cadherin cluster lifespan distribution of control and Xdd1 clusters. Conditions were statistically compared using a Kolmogorov-Smirnov test.

## Discussion

This work is an investigation into the relationship between PCP and cadherin-based cell adhesions during *Xenopus* gastrulation. Our initial question asked if Cdh3 was planar polarized to t-junctions during *Xenopus* CE, as would be expected based on results from *Drosophila* germband CE (Blankenship et al., 2006). However, we found that Cdh3 became increasingly enriched at shortening v-junctions. This result hints at a critical difference in the behavior of cadherins during *Xenopus* CE compared to Drosophila. Next, we found that PCP was required for coordinated oscillations of Cdh3 and actin, suggesting PCP controls the coupling of cadherin and actin dynamics. This result was also of interest because previous perturbations of actin or cadherin oscillations altered the frequency (Shindo et al., 2019) or amplitude (Huebner et al., 2022) of oscillations. However, Xdd1 only controlled the coupling of the actin and cadherin dynamics but did not appear to alter the amplitude or frequency of the oscillations.

Finally, we wanted to investigate the relationship between PCP and cadherin clustering as two previous reports linked PCP to clusters in Xenopus (Huebner et al., 2021; Kraft et al., 2012). To this end we developed a method to measure the size, number, and lifespan of individual cadherin clusters. Previous research has shown cadherins physically interacting with the PCP proteins Vangl and Wnt-11/Frizzled-7, and in both cases the interaction with PCP components is thought to sequester cadherins (Kraft et al., 2012; Nagaoka et al., 2014). Here we perturbed the core PCP protein Dishevelled and found that this perturbation resulted in smaller and shorter-lived cadherin clusters. This result indicates another interaction between cadherins and a core PCP protein, dishevelled-2. Further we show for the first time that PCP has a role in controlling the stability of cadherin clusters. While there is still little known about the relationship between PCP and cadherins here we show a clear connection between PCP and the ability to form stable cadherin clusters and this work should provide a foundation for further investigation into the connection between PCP and cadherins.

## Material and methods

### *Xenopus* embryo manipulations

Adult female *Xenopus* were induced to ovulate by injection of 600 units of human chorionic gonadotropin and animals were kept at 16°C overnight. The following day the ovulating females were gently squeezed to stimulate egg laying and then eggs were fertilized in vitro. Approximately 1.5 hours after fertilization embryos were dejellied with 3% cysteine (pH 8) for 10 minutes and then washed in 1/3X Marc’s Modified Ringer’s (MMR) solution. Prior to microinjection embryos were placed in 2% ficoll in 1/3X MMR and following microinjection embryos were reared in 1/3X MMR. Embryos were injected using a Parker’s Picospritizer III with an MK1 Manipulator. Embryos were injected at the 4-cell stage in the dorsal blastomeres targeting the presumptive dorsal marginal zone. Embryos were dissected at stage 10.25 in Steinberg’s solution to isolate Keller explants (Keller et al., 1992).

### Plasmids, antibody, and morpholino

Cdh3-GFP (Pfister et al., 2016), Cdh3-cis-mutant (Huebner et al., 2021), lifeact-RFP, and membrane-BFP plasmids were made in pCS105 and Xdd1 (Sokol, 1996) was made in CS2myc. The Cdh3 antibody was from the Developmental Studies Hybridoma Bank (catalog number 6B6). The Cdh3 morpholino was ordered from Gene Tools and had been previously characterized (Ninomiya et al., 2012).

### mRNA microinjections

Capped mRNAs were generated using the ThermoFisher SP6 mMessage mMachine kit (catalog number AM1340). mRNAs were injected at the following concentration per blastomere, Cdh3-GFP (50pg), Cdh3-cis-mutant (300 pg), lifeact-RFP (100pg), membrane-BFP (100pg), and Xdd1 (1ng). Cdh3 morpholino was injected at a concertation of 10ng per blastomere.

### Imaging *Xenopus* explants

Following dissection explants were incubated at room temperature for 4 hours or 16°C overnight prior to imaging. Explant were maintained in either Steinberg’s solution or Danilchik’s for Amy solution during imaging and were mounted on fibronectin coated glass coverslips. Super-resolution images were acquired with a BioVision Technologies instantaneous structured illumination microscope. Standard resolution confocal images were acquired with either a Zeiss LSM 700 or a Nikon A1R. Super-resolution movies were collected with a 2 second time interval and confocal movies were acquired with a 20 second time interval. All images were acquired at an approximate depth of 5μm into the tissue.

### Measurement of protein intensities at cell-cell junctions

The open-source image analysis software Fiji (Schindelin et al., 2012) was used for image processing and quantification. First, to better visualize the cadherin clusters, images were processed with a 50-pixel rolling ball radius background subtraction and smoothed with a 3×3 averaging filter. The segmented line tool was then used to set a line of interest across the length of the cell-cell junction with a line with width set to the thickness of the junction (16 pixels). The measure tool was then used to measure the mean intensity values. For time lapse movies we used the Fiji time lapse plugin line interpolator tool to make successive measurements for each timepoint. Here a LOI was drawn every 10-30 frames and the line interpolator tool was used to fill in the LOIs between the manually drawn in LOIs. The measure tool was then used to extract mean intensities at each timepoint.

### Measurement of cadherin cluster sizes and lifespans

Here we used the Fiji time lapse line interpolator tool to draw LOIs (as described above) across the length of a cell junction for 200 frames (400 seconds) of a movie. Then the time lapse line interpolator tool was used to generate a kymograph of all the LOIs. Kymographs were thresholded at 2 times the mean intensity of the kymograph. We then used a 40-pixel smooth along the time axis to connect the LOIs. The Fiji analyze particle tool was used to identify individual clusters and clusters that hit the edge of the image were excluded so that we only analyzed complete clusters. We next used the measure bounding rectangle tool to identify the width (cluster size) and length (cluster lifespan) of each cluster. Finally, clusters of 1-2 pixels in width were filtered out as noise.

### Fluorescent recovery after photobleaching

Bleaching experiments were acquired on a Zeiss 700 confocal with a 4 second time interval and images were acquired for four minutes post bleaching. Regions of interest were bleached using 405nm and 488nm wavelength lasers set at 35% power. A second region of interest was from a neighboring non-bleached cell was used for bleach correction. Bleach correction and curve-fitting was carried out using a Python script (modified from https://imagej.net/tutorials/analyze-frap-movies).

### Cdh3 immunostaining

Explants were prepared as described above and then fixed in 1x MEMFA for one hour at room temperature. Samples were then washed three times in PBS to remove fixative and permeabilized with 0.05% Triton X-100 in PBS for 30 minutes. Next samples were blocked in 1% normal goat serum (NGS) in PBS for two hours at room temperature. The primary antibody was diluted 1:100 in 1% NGS/PBS and samples were incubated with primary antibody at 4°C overnight. We then performed a second incubation in blocking solution for one hour at room temperature. Secondary antibody (goat anti-mouse 488, #A32723) was diluted 1:500 and samples were incubated with secondary antibody for 1 hour at room temperature. Finally, samples were washed three times in 1x PBS and imaged

## Supplemental figure legends

**Supplemental Figure 1:**
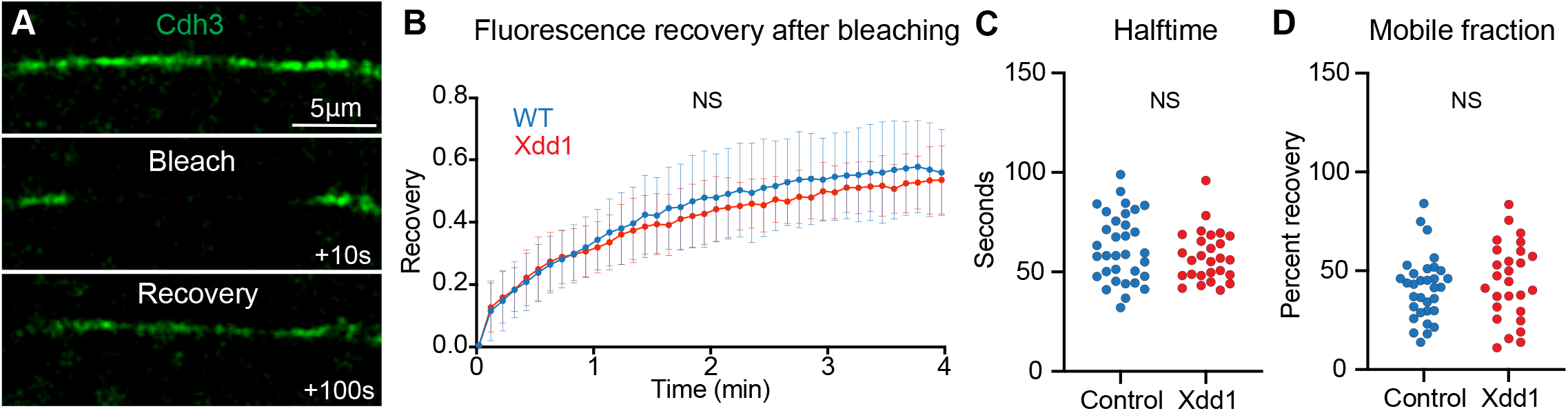
Cdh3 turnover is not changed in the presence of Xdd1. **A**. Time series of a single junction labeled with Cdh3 that is bleached and then allowed to recover. **B**. Graph showing the bleach recovery for Cdh3 in control cells or cells expressing Xdd1. Conditions were statistically compared using a Kolmogorov-Smirnov test and were not significantly different. **C**. Graph comparing the recovery halftime of Cdh3 for control or Xdd1 expressing cells. Conditions were not significantly different as assessed by a Mann Whitney test. **D**. Graph comparing the mobile fraction of Cdh3 for control and Xdd1 expressing cells. Conditions were compared using a Mann Whitney test and there was no significant difference.

**Supplemental figure 2:**
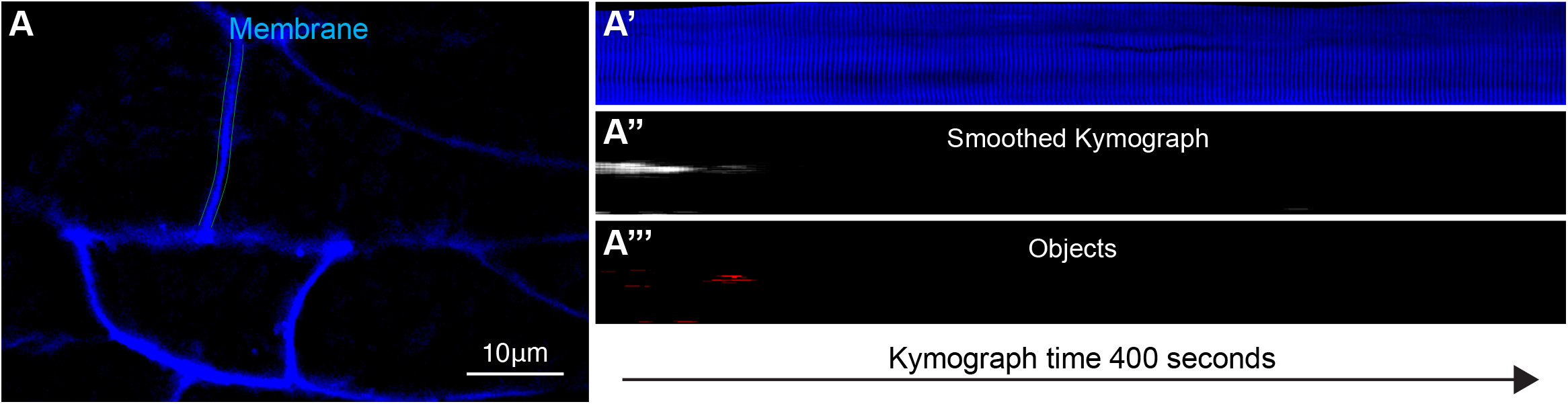
Membrane marker as a negative control for cluster analysis. **A**. Still frame from a time lapse movie labeled with a membrane marker shown in blue. Teal lines highlight the junction that is displayed as a kymograph in Supp. Fig.1A’. **A’**. Kymograph showing the membrane marker over a 400 second time interval. **A’’**. Time smoothed kymograph of the image shown in Supp. Fig.1A’ **A’’’**. Kymograph of the membrane marker after thresholding and converting to objects.

**Supplemental figure 3:**
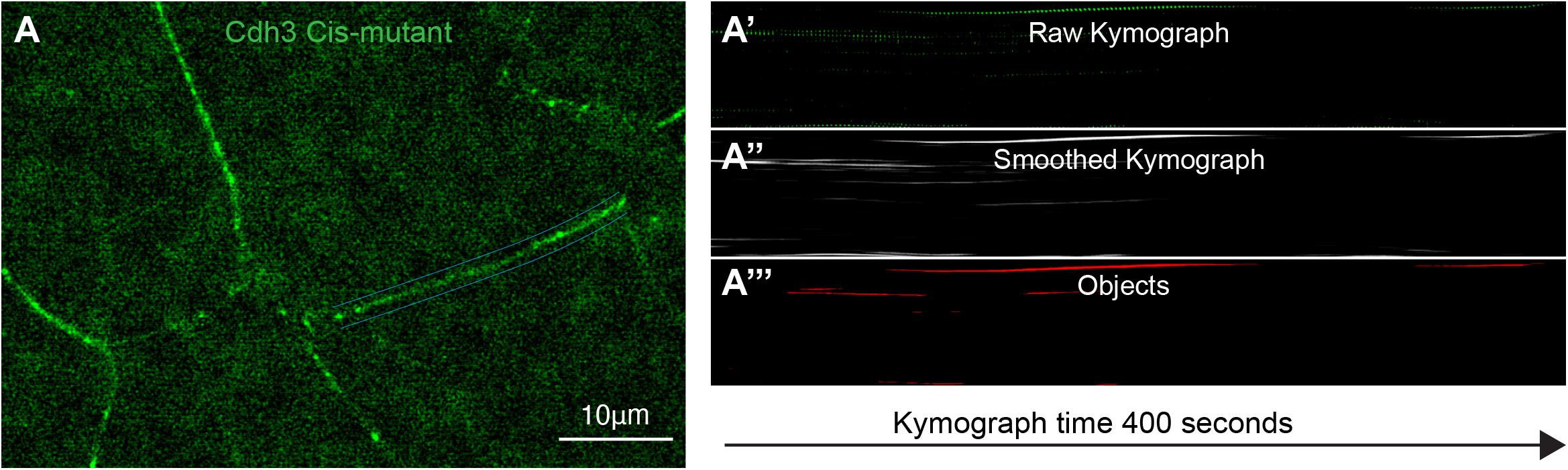
Cdh3 cis-mutant forms fewer and smaller clusters. **A**. Still frame from a time lapse movie of the Cdh3-cis-mutant shown in green. Blue lines highlight the junction that is displayed as a kymograph in Supp. Fig.2A’. **A’**. Kymograph showing the Cdh3-cis-mutant over a 400 second time interval. **A’’**. Time smoothed kymograph of the image shown in Supp. Fig.2A’ **A’’’**. Kymograph of the Cdh3-cis-mutant after thresholding and converting to objects.

